## Supplementary Table 4 for "A single amino acid polymorphism in a conserved effector of the multihost blast fungus pathogen expands host-target binding spectrum"

**Data Collection and Refinement Table**

|  | **APikL2A / sHMA25** | **APikL2F / sHMA94** |
| --- | --- | --- |
| **Data collection statistics** |  |  |
| Wavelength (Å) | 0.9795 | 0.9119 |
| Space group | P 21 21 21 | C 2 2 21 |
| Cell dimensions | 46.2 46.7 89.0 | 125.4 222.5 102.0 |
| *a*, *b*, *c* (Å) | 90.0 90.0 90.0 | 90.0 90.0 90.0 |
| Resolution (Å)* | 41.0-1.8 (1.84 – 1.80) | 109.51-2.3 (2.36-2.30) |
| *R*_meas_ (%) | 5.8 (141.9) | 8.7 (117.4) |
| *I*/σ*I* | 21.5 (1.8) | 10.6 (1.4) |
| Completeness (%) |  |  |
| Overall | 100 (100) | 99.9 (99.8) |
| Anomalous | 100 (100) | 99.5 (99.6) |
| Unique reflections | 18505 (1060) | 63610 (4432) |
| Redundancy |  |  |
| Overall | 13.0 (12.7) | 6.6 (6.6) |
| Anomalous | 6.9 (6.6) | 3.3 (3.3) |
| CC(1/2) (%) | 99.9 (79.8) | 99.9 (75.6) |
| **Refinement and model statistics** |  |  |
| Resolution (Å) | 41.0-1.8 (1.84 – 1.80) | 109.5-2.3 (2.36-2.30) |
| *R*_work/_*R*_free_ (%) | 19.6 / 23.5 | 20.9 / 24.6 |
| No. atoms |  |  |
| Protein | 2677 | 10600 |
| Ligand | 20 | 111 |
| B-factors |  |  |
| Protein | 47.58 | 61.92 |
| Ligand | 42.8 | 74.26 |
| R.m.s deviations |  |  |
| Bond lengths (Å) | 0.0147 | 0.0152 |
| Bond angles (º) | 1.92 | 1.92 |
| Ramachandran plot (%)** |  |  |
| Favoured | 96.97 | 97.88 |
| Allowed | 2.42 | 2.12 |
| Outliers | 0.61 | 0 |
| MolProbity Score | 1.62 |  |

*The highest resolution shell is shown in parenthesis.

**As calculated by MolProbity
